# Enhanced representation of space by prefrontal neuronal ensembles and its dependence on cognitive states

**DOI:** 10.1101/065581

**Authors:** Mohammad-Reza A. Dehaqani, Abdol-Hossein Vahabie, Mohammadbagher Parsa, Behard Noudoost, Alireza Soltani

**Affiliations:** School of Cognitive Sciences, Institute for Research in Fundamental Sciences, Tehran, Iran.; Department of Computer Science, Montana State University, Bozeman MT; Department of Cell Biology and Neuroscience, Montana State University, Bozeman MT; Department of Psychological and Brain Sciences, Dartmouth College, Hanover NH 03755

## Abstract

Although individual neurons can be highly selective to particular stimuli and certain upcoming actions, they can provide a complex representation of stimuli and actions at the level of population. The ability to dynamically allocate neural resources is crucial for cognitive flexibility. However, it is unclear whether cognitive flexibility emerges from changes in activity at the level of individual neurons, population, or both. By applying a combination of decoding and encoding methods to simultaneously recorded neural data, we show that while maintaining their stimulus selectivity, neurons in prefrontal cortex alter their correlated activity during various cognitive states, resulting in an enhanced representation of visual space. During a task with various cognitive states, individual prefrontal neurons maintained their limited spatial sensitivity between visual encoding and saccadic target selection whereas the population selectively improved its encoding of spatial locations far from the neurons' preferred locations. This 'encoding expansion' relied on high-dimensional neural representations and was accompanied by selective reductions in noise correlation for non-preferred locations. Our results demonstrate that through recruitment of less-informative neurons and reductions of noise correlation in their activity, the representation of space by neuronal ensembles can be dynamically enhanced, and suggest that cognitive flexibility is mainly achieved by changes in neural representation at the level of population of prefrontal neurons rather than individual neurons.

## Introduction

An ensemble of neurons could provide a complex representation of external stimuli, ongoing processes, or upcoming actions based on multiplexed response of neurons contained in that ensemble [1–3]. At the same time, individual neurons could be highly selective to external stimuli [4–6] or highly informative about the animal's action [7–9] or performance [10]. These beg the question of why downstream neurons should read out/rely on the activity of neural ensembles instead of the activity of selective/informative individual neurons. Naturally, neural representation based on population activity could improve the information content compared to individual neurons, but are there additional advantages for population coding beyond such improvement?

Recently, the combination of improved techniques for large-scale recording from neuronal ensembles and novel computational methods has led to a surge of interest in understanding the contribution of population-level representation to cognitive functions such as working memory, visual attention, decision making, and categorization [1,3,11–14]. On the one hand, a large body of theoretical work [15–21] and electrophysiological studies [22–27] has revealed the contribution of inter-neuronal correlations to the coding capacity of neural populations. For example, it has been shown that noise correlation affects signal-to-noise ratio of the population response [22] and modulates sensory coding more than the firing rate of individual neurons [25,26]. On the other hand, others have argued that high-dimensional neural representations are more valuable for cognitive functions than representations based on highly specialized neurons [1,3,28,29]. Perhaps, which of these factors (or both) is crucial could reveal mechanisms by which a given population contributes to a cognitive function. For example, enhanced encoding due to a decrease in noise correlation could indicate that the population activity is modulated by an external source (e.g. attentional modulation) whereas high-dimensional representation could point to interactions between neurons in the same population.

We hypothesized that enhancement of the information content of a population of neurons beyond its individual neurons is dynamically adjusted by cognitive states. To test this hypothesis, we simultaneously recorded neural activity in the Frontal Eye Field (FEF) of monkeys during an oculomotor delayed response task. This task requires encoding of the location of a target stimulus and making a saccade to the remembered location after a delay and thus, involves visual encoding, maintenance of spatial information, and saccadic target selection. Employing a combination of encoding and decoding methods to simultaneously recorded neural activity, we found a selective enhancement of the information content of the population of neurons prior to saccades. More specifically, although individual neurons maintained their limited spatial sensitivity from visual encoding to saccadic target selection, the ability of the population to encode spatial locations far from the neurons' preferred locations improved, a phenomenon which we refer to as 'encoding expansion'. In other words, locations poorly encoded by individual neurons (non-preferred locations) could be discriminated much better at the population level. This enhanced population-level information relied on high-dimensional neural representations for non-preferred locations and was accompanied by selective reductions in noise correlation for the same locations. Overall, we show for the first time that by differently impacting preferred and non-preferred locations of visual space, cognitive states can dynamically enhance the population-level information while keeping the individual neuronal selectivity rather unchanged.

## Results

Using 16-channel linear array electrodes (V-probe, Plexon Inc.), we simultaneously recorded from a population of neurons in the FEF of two monkeys during an oculomotor delayed response (ODR) task (188 single neurons, 1165 pairs of neurons; see Materials and methods). The ODR task (Fig 1A) enabled us to trace the transformation of visual input into motor action (saccade), and has been classically used to determine the contribution of FEF neurons to each of these processes [30–32]. Our goal was to determine how visual space is encoded differently by a given ensemble of neurons and the participating individual neurons, and how this differential encoding of space depends on two cognitive states (visual encoding and saccadic target selection). Therefore, we analyzed the simultaneously recorded neural data to measure spatial information encoded at the single-cell and population levels, as well as information embedded in the correlated activity of pairs of neurons.

**Fig 1.**
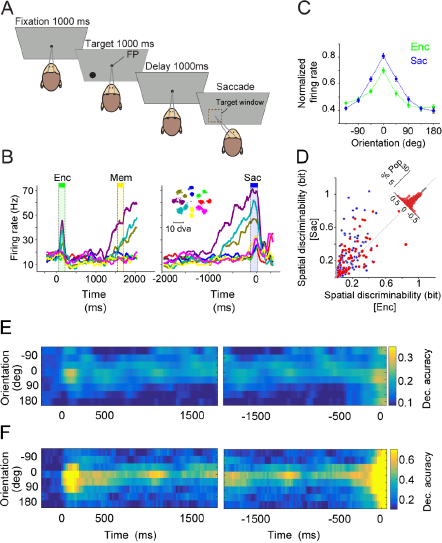
Spatial encoding of a population of FEF neurons expands beyond the ability of individual neurons prior to saccade. **(A**) Behavioral paradigm. Each trial of the oculomotor delayed response task started with fixation on a central spot for 1000 msec followed by the presentation of a visual target (black circle) for 1000 msec in one of the 16 possible locations (at 8 angles and 2 eccentricities). Disappearance of the target was followed by a 1000 msec delay, after which the central spot disappeared and the monkey made a saccade to the remembered target location. (**B**) The mean response of an example FEF neuron to the target presented at 8 different angles (averaged over two eccentricities), indicated by color in the inset. The neural response is aligned to the onset of the target (left panels) or saccade (right panel). (**C**) Normalized response of 188 FEF neurons to targets presented at different angles relative to the neuron's preferred location during visual encoding (Enc, green) and saccade preparation (Sac, blue) epochs. The error bars represent s.e.m. The spatial tuning of individual neurons only slightly improved from visual encoding to saccade preparation. (**D**) Comparison of the spatial discriminability during visual encoding and saccade preparation. Histogram on the diagonal shows the difference in spatial discriminability, and FEF neurons exhibiting mixed selectivity are shown in red (*n* = 65). (**E-F**) Plotted is the decoding accuracy for different target locations based on the SVM classifier, when the activity of individual units (E) or the ensemble of units (F) recorded in an example session is used. The time course of decoding accuracy showed similar accuracy during saccade preparation and visual encoding for individual units (E) but larger accuracy by the ensemble of neurons during saccade preparation than visual encoding for nonpreferred locations. The left (right) panels shows the results when the activity is aligned to target (saccade) onset.

### Presaccadic expansion of the population code

The average activity of an example recorded neuron during the ODR task revealed the typical spatial tuning/selectivity of FEF neurons (Fig 1B). This neuron responded differently to stimuli presented at eight possible angles during both visual encoding (Enc, one way Anova, F = 18.51, *p* < 10^−3^) and just before the saccade to the remembered location (Sac, one way Anova F = 48.72, *p* < 10^−3^). However, the neural response was more graded during saccade preparation than visual encoding. Similarly, the average normalized activity across all recorded neurons showed stronger spatial tuning during saccade preparation compared to visual encoding such that the tuning was narrower for the former epoch (tuning dispersion: Sac = 2.06±0.04, Enc = 2.25±0.04, *p* < 10^−3^; see Materials and methods). We found a similar relationship between the response during saccade preparation and visual encoding when we restricted the analysis to the population of neurons that showed selectivity in more than one task epoch (i.e. mixed selectivity neurons, *n* = 65, S1 Fig).

We next quantified the information content of individual neurons using the mutual information between the response and stimulus location (see Materials and methods). We refer to this mutual information as 'spatial discriminability' as it quantifies the ability of individual neurons to discriminate between various target locations. For the example neuron, the spatial discriminability was greater during saccade preparation than visual encoding (spatial discriminability: Enc = 0.09±0.01, Sac = 0.18±0.02, *p* < 10^−3^). This difference was also observed across all neurons such that the spatial discriminability was greater during saccade preparation than visual encoding for the entire population of neurons as well as selective neurons (Enc_all_ = 0.15±0.01, Sac_all_ = 0.23±0.02, *p* < 10^−3^; *n* = 188, Enc_selective_ = 0.21±0.02, Sac_selective_ = 0.27±0.02, *p* = 0.007; Fig 1D). This enhanced representation of saccade targets is consistent with previous studies [30] and the known role of FEF in oculomotor control.

In addition to mutual information, which reflects the spatial discriminability of single neurons, we employed a support vector machine (SVM) to quantify the ability of the population of FEF neurons and individual FEF neurons to represent the target location. The decoding accuracy measures the ability of an external observer (the classifier) to determine the location of the target solely based on the simultaneous activity of recorded single units and multi units. The SVM classifier was designed to classify eight target angles either when all the units were considered individually (i.e. separately running the SVM for each unit, SVM_individual_) or when the simultaneous activity of the ensemble of units recorded in one session was taken into account (i.e. running the SVM for each recording session, SVM_ensemble_; see Materials and methods). The time course of the SVM performance applied to 36 individual units recorded within a single recording session of the experiment showed similar accuracy during saccade preparation and visual encoding (Fig 1E). However, when the activity of the population of simultaneously recorded units was taken into account, the decoding accuracy during saccade preparation improved dramatically (Fig 1F). This improvement was most pronounced for locations in the opposite hemifield. For example, the population decoding accuracy for the 180-degrees away location was about 5 times larger than the decoding accuracy of individual units (decoding accuracy_ensemble_= 0.6±0.09, considering the chance level of 0.125). In contrast, the decoding accuracy for individual units was 0.34±0.18 for the preferred location and dropped to only 0.23±0.22 for a location 180 degrees away. Altogether, results from this example recording session revealed a population-level enhancement of neural encoding during saccade preparation ('presaccadic expansion') which occurred primarily for parts of space weakly encoded by the single-cell response.

The presaccadic expansion of the neural code was consistently observed across all recording sessions. For each unit or session, we determined the preferred location based on the decoding accuracy of the SVM taking the whole visual, memory and presaccadic activity into account. Overall, we found that preferred locations showed almost no presaccadic change in decoding accuracy at the level of individual units (Δdecoding accuracy_individual_ = 0.0074±0.0135, *p* = 0.0.722, *n* = 481; Fig 2A). At the same time, nonpreferred locations (>90 degrees away from the preferred location) exhibited a modest change (Δdecoding accuracy_individual_ = 0.0313±0.0042, *p* < 10^−3^). In contrast, when the activity of the population of simultaneously recorded units was taken into account, the decoding accuracy during the presaccadic period improved dramatically for both preferred and nonpreferred locations. This presaccadic enhancement was observed across all eight locations, as indicated by the greater overall decoding accuracy in SVM_ensemble_ during saccade preparation than visual encoding (Δdecoding accuracy_ensemble_ = 0.09±0.03, *p* = 0.008; Fig 2A). Importantly, this presaccadic population benefit was not equally strong across all target locations: the benefit was small for the neurons' preferred location (Δdecoding accuracy_pref_ = 0.02±0.03; *p* = 0.711), and much larger for locations far from the preferred locations (Δdecoding accuracy_nonpref_ = 0.12±0.03, *p* = 0.008; Fig 2B).

**Fig 2.**
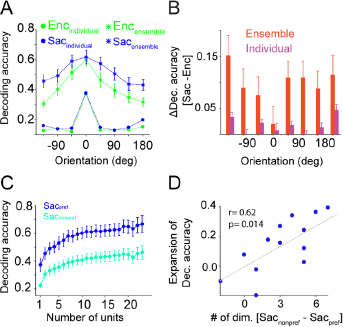
Presaccadic expansion relies on high-dimensional neural representation of space. (**A**) Plotted is the decoding accuracy (Dec. accuracy) of the SVM classifier for various locations (relative to preferred) for individual (circles) and ensemble (stars) of units during visual encoding (Enc) and saccade preparation (Sac). The decoding accuracy of the ensemble of units for nonpreferred locations remarkably increased from Enc to Sac. (**B**) The change in decoding accuracy between visual encoding and saccade preparation for individual units and the ensemble of units as a function of target locations. Larger improvements are visible for nonpreferred locations. (**C**) The decoding accuracy of the ensemble of units plotted as a function of the number of units, separately for the preferred and nonpreferred target locations during saccade preparation. (**D**) The difference in dimensionality for preferred and nonpreferred locations predicted encoding expansion. Plotted is the encoding expansion index during a given session as a function of the difference in dimensionality between nonpreferred and preferred locations during Sac in that session. The dashed line illustrates least-squares line fitted to points and the error bars represent s.e.m.

To rule out the possibility that the observed encoding expansion is merely due to engagement of neurons with certain types of selectivity during different task epochs (e.g. visual neurons during visual encoding and motor neurons during saccade preparation), we computed changes in the decoding accuracy of the SVM classifier for different types of neurons (S2 Fig A-B). We observed similar encoding expansion for the visual, motor, and visuomotor neurons. For visual neurons changes in decoding accuracy were:Δdecoding accuracy_pref_ = −0.03±0.03, *p* = 0.376; Δdecoding accuracy_nonpref_ = 0.09±0.03, *p* = 0.027. For motor neurons these changes were: Δdecoding accuracy_pref_ = 0.03±0.03, *p* = 0.494; Δdecoding accuracy_nonpref_ = 0.12±0.03, *p* = 0.003. Finally, for visuomotor neurons changes in the decoding accuracy were: Δdecoding accuracy_pref_ = −0.05±0.03, *p* = 0.1; Δdecoding accuracy_nonpref_ = 0.1±0.03, *p* = 0.005 (S2 Fig C-E). These results indicate that observed encoding expansion is not due to particular contributions of different types of neurons during different task epochs.

Altogether, these results reveal an expansion of the population code prior to saccadic target selection. Such an expansion can serve as a potential mechanism to enhance the robustness or precision of the population code by incorporating a larger ensemble of FEF neurons. Note that this presaccadic enhancement was mainly observed at the population level and not the individual units (Fig 2B), implying that the expansion of the neural code relies on high-dimensional neural representations and also depends on changes in the inter-neuronal relationships. Thus, we next examined the contributions of these factors to presaccadic expansion.

### Presaccadic expansion relies on high-dimensional neural representations

We found that during saccade preparation, the ability of the population to represent target locations expanded far beyond the capability of single neurons: locations which were poorly encoded by single neurons could be accurately discriminated using an ensemble-level population read-out. Recent studies have indicated that high-dimensional neural representations are beneficial for information encoding [1,3]. Therefore, we next examined whether the encoding expansion during saccade preparation was associated with a greater dimensionality in the population.

To do so, we sorted the simultaneously recorded units in each session based on their mutual information (spatial discriminability), and made a set of neural ensembles by subsequently adding new units (with decreasing mutual information). Using this approach, we first computed the overall decoding accuracy as a function of the number of units (NU) in different epochs of the experiment (S3 Fig). We found that about 4 units are enough to explain 75% of the ultimate classifier performance during both saccade preparation and visual encoding (NU_Enc_ = 4.44±0.84, NU_Sac_ = 3.82±0.67, errors based on bootstrapping; see Materials and methods).

The encoding expansion during saccade preparation occurred because the SVM performance increased more strongly for nonpreferred than preferred locations. Thus, we computed the dimensionality of neural representation separately for preferred and nonpreferred (>90 deg) locations (Fig 2C). We found that 6.54±2.21 units were necessary to achieve 75% of the whole-ensemble level of performance for nonpreferred locations, compared to only 2.22±0.45 units for preferred locations, indicating that the population response had a much larger dimensionality for representing nonpreferred compared to preferred locations (bootstrap confidence interval, *p* < 10^−3^; Fig 2C). We obtained qualitatively similar results when we performed dimensionality analysis for neurons with different types of selectivity (visual neurons: NU_pref_ = 4.16±0.9, NU_nonpref_ = 10.46±3.64, *p* < 10^−3^; motor neurons: NU_pref_ = 2.76±0.73, NU_nonpref_ = 9.38±2.71, *p* < 10^−3^; visuomotor neurons: NU_pref_ = 4.89±1.05; *p* = 0.376, NU_nonpref_ = 10.76±2.54, *p* < 10^−3^; *p*-values are computed by bootstrap confidence interval; S2 Fig F-H). Note that the exact value of the dimensionality strongly depends on the threshold (here 75%) such that a larger dimensionality is obtained with a larger value of the threshold. Nevertheless, our results indicate that about three times more neurons are needed to encode nonpreferred locations than preferred locations.

We next examined the direct relationship between the encoding expansion and the difference in dimensionality between nonpreferred and preferred locations by defining an expansion index for each recording session of the experiment (see Materials and methods). There was a positive correlation between the encoding expansion index and the difference in dimensionality between nonpreferred and preferred locations (Spearman correlation, *r* = 0.62, *p* = 0.014; Fig 2D). This demonstrates that a larger difference in dimensionality between nonpreferred and preferred locations is accompanied by a larger increase in the population-level, differential encoding of nonpreferred than preferred locations during saccade preparation (i.e. presaccadic expansion). These results suggest that high-dimensional neural representations are crucial for the observed presaccadic expansion.

### Changes in inter-neuronal correlations contribute to encoding expansion during saccade preparation

As mentioned above, reduction in noise correlation has been proposed as a mechanism to enhance population-level information [15,18,20,22,26]. Hence, we examined the effects of inter-neuronal correlations at the level of pairs of neurons (see Materials and methods) in order to address the contribution of correlated neural activity to the enhanced spatial representation prior to saccades. The correlated activity of a pair of neurons could be decomposed into two components: 1) signal correlation (SC), which in this task reflects the degree to which two neurons exhibit similar response to a given target; and 2) noise correlation (NC), measuring shared variability unrelated to the target location [33]. By transforming the problem of population level analysis to the level of pairs of neurons and stimuli (~ 32400 neuron-stimulus pairs), we were able to precisely study the contribution of noise and signal correlations to encoding expansion during saccade preparation.

We found a higher SC during saccade preparation than visual encoding (SC_Enc_ = 0.0164±0.0005, SC_Sac_ = 0.0447±0.0008, *p* < 10^−3^). This greater correlation in the presaccadic spatial tuning of FEF neurons was consistent with the enhanced response (Fig 1C) and spatial discriminability (Fig 2A) observed during this epoch within the same population. In contrast to SC, the NC was reduced in the presaccadic period (NC_Enc_ = 0.046±0.0007, NC_Sac_ = 0.039±0.0007, *p* < 10^−3^; Fig 3A). Interestingly, the presaccadic reduction in NC was prominent for pairs of nonpreferred locations (>90 deg) compared to preferred locations (ΔNC_nonpref_ = -0.0152±0.0029, ΔNC_pref_ = -0.0007±0.0025, *p* < 10^−3^; Fig 3A); however, this differential change was not observed for increases in signal correlation (ΔSC_pref_ = 0.0139±0.0021, ΔSC_nonpref_ = 0.0104±0.0012, *p* = 0.376). To precisely measure the modulation of noise correlation during Sac compared to Enc, we assessed the tuning of noise correlation across Sac and Enc. For locations far from the preferred location, NC changed strongly during Sac compared to Enc interval (Fig 3B). This differential change in noise correlation (from visual encoding to saccade preparation) across space may provide a population-level advantage for processing of nonpreferred locations. We also confirmed that this differential decrease in noise correlation was present for all neuron types (S4 Fig A-C) and dovetailed with the presaccadic expansion of population code measured by the SVM.

**Fig 3.**
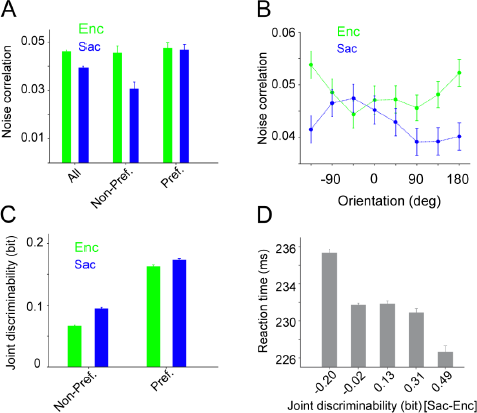
Noise correlation and information content of pairs of neurons (joint discriminability) differentially change for preferred and nonpreferred locations, and the behavioral relevance of the latter. (**A**) Average noise correlation across all locations and separately for preferred and non-preferred locations during visual encoding (Enc, green) and saccade preparation (Sac, blue). There was an overall reduction in noise correlation during Sac, but this effect was driven by a reduction in noise correlation at nonpreferred locations. (**B**) Selective reduction in noise correlation for non-preferred locations during saccade preparations. Plotted is the tuning curve of noise correlation during visual encoding and saccade preparation. (**C**) Plotted is the joint discriminability separately measured for preferred and nonpreferred locations during visual encoding and saccade preparation. Nonpreferred locations showed a larger improvement in joint discriminability from visual encoding to saccade preparation. (**D**) Faster reaction times corresponded with greater improvement in joint discriminability from visual encoding to saccade preparation. The error bars represent s.e.m.

Because signal and noise correlations are defined based on the activity of pairs of neurons and not the whole populations, computing their contribution to the observed encoding expansion of the population code requires measuring the information content of pairs of neurons (as a proxy for the population). Thus, we employed a different approach to reduce the study of population to the level of pairs of neurons. More specifically, we measured the information content of a given pair of neurons by the ability of this pair to discriminate a pair of locations ('joint' discriminability) using the linear discriminant analysis (LDA) technique (see Materials and methods and S5 Fig A). Such approach enables us to precisely measure the contribution of NC and SC to the enhancement of neural representation. Consistent with the SVM results, joint discriminability was enhanced during saccade preparation compared to visual encoding (Enc = 0.1572±0.0009, Sac = 0.2274±0.0013, *p* < 10^−3^). Moreover, similar to the SVM results (Fig 2A), nonpreferred locations showed greater presaccadic improvements in discriminability than preferred locations (Δjoint discriminability_nofpref_ = 0.0281±0.0027, Δjoint discriminability_pref_ = 0.0105±0.0031, *p* < 10^−3^; Fig 3C), and qualitatively similar results were obtained for all neuron types (S4 Fig D-F).

Considering similarities between changes in joint discriminability and decoding performance, we next tested whether the differential improvement in joint discriminability for nonpreferred than preferred locations reflects the encoding expansion measured by the SVM. To do so, we computed the correlation between the encoding expansion index using the SVM and a similar index based on joint discriminability after subtracting a baseline related to the information content of individual cells (see Materials and methods). A significant correlation between encoding expansion index based on the SVM and joint discriminability demonstrated that joint discriminability reflects encoding expansion (Spearman correlation, *r =* 0.53 *p =* 0.025). Overall, the analyses reported above based on pairwise measures confirmed the expanded presaccadic spatial encoding originally revealed by the decoding analysis, and demonstrated that this expansion is traceable in the activity of as few as two neurons.

Unlike encoding expansion based on the SVM, joint discriminability measures the information content of a pair of neurons on a trial-by-trial basis and thus, can be used to look for behavioral correlates of changes in the information content between visual encoding and saccade preparation. By binning the change in joint discriminability from visual encoding to saccade preparation, we found that greater changes in joint discriminability were associated with faster reaction times (Fig 3D). More specifically, there was ~5% difference between the average reaction time when the joint discriminability was negative and positive (235.32±0.39 vs. 224.60±0.68 msec, *p* < 10^−3^), showing that changes in joint discriminability had a significant impact on the behavior.

Having established that changes in joint discriminability reflect encoding expansion, we next examined the relationship between signal correlation, noise correlation, and joint discriminability for the same pairs of neurons during visual encoding and saccade preparation. This analysis allowed us to study the contribution of signal and noise correlations to encoding expansion. To do so, we first partitioned our dataset according to the sign of signal and noise correlations in each pair. Interestingly, we found that indeed when two neurons had a common source of signal (a positive SC), they were three times more likely to have a common source of noise (a positive NC; S6 Fig). Moreover, pairs with a positive signal and noise correlations comprised about 45% of the dataset. Finally, presaccadic expansion based on joint discriminability was at least two times greater in these pairs than in those pairs with one or more negative correlation values (S6 Fig).

By examining the relationship between signal correlation and joint discriminability during visual encoding for pairs of neuron with positive signal and noise correlations, we found that a higher signal correlation was associated with greater discriminability (Fig 4A). However, by separating the population based on whether noise correlation decreased or increased during the presaccadic period (comprising 76% and 24% of population, respectively), we found that for a given level of SC the joint discriminability depended upon the change in noise correlation such that a decrease in noise correlation improved joint discriminability. This result indicated that a change in noise correlation was sufficient to improve encoding expansion during saccade preparation (also see S7 Fig A).

**Fig 4.**
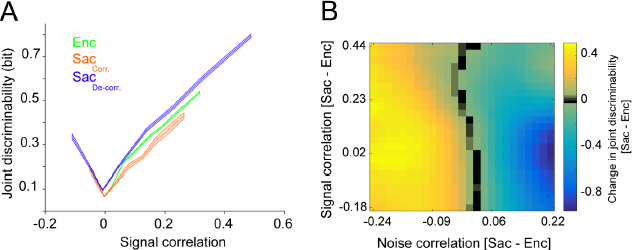
Changes in noise correlation contribute to presaccadic expansion independent of signal correlation. (**A**) Joint discriminability depends upon both signal and noise correlation. Plotted is joint discriminability as a function of signal correlation computed separately for neuron-location pairs during visual encoding (Enc, green) and during saccade preparation. The latter is plotted for pairs of neurons that showed a large (> 0.1) reduction (blue) or increase (red) in noise correlation from visual encoding to saccade preparation. The shaded area represents s.e.m. For a given value of signal correlation, joint discriminability of neurons that became decorrelated was larger than those which became more correlated during saccade preparation. (**B**) Change in joint discriminability was explained by a reduction in noise correlation when the change in signal correlation was taken into account. The change in joint discriminability (separately z-scored across the same change of signal correlation, each row) is plotted as a function of the changes in signal and noise correlations.

As a further control, we verified that the isolated discriminability (which excludes the contribution of NC yet incorporates each unit's spatial sensitivity) stayed the same between neurons for which NC decreased during saccadic preparation and those for which NC increased (S7 Fig B). Finally, we examined the presaccadic change in joint discriminability as a function of changes in SC and NC and found that the change in discriminability depended on the change in NC, independent of the change in SC (Fig 4B). Overall, these results indicated that decorrelation (i.e. a reduction in noise correlation) contributed to the presaccadic encoding expansion.

### Discussion

The brain's processing capacity is limited by physical constraints including intrinsic noise in neural firing, the number of neurons, and their connectivity. Therefore, changes in neural encoding which improve the ability of a fixed population of neurons to represent information are of particular interest to neuroscientists, because such changes enable the brain to dynamically allocate limited neural resources based on task demands and to maximize neural computations [34,35]. Here, we examined neural representations of saccade targets in the FEF and found dynamic improvement in spatial encoding according to cognitive states at the population level, but specifically for parts of space not well-encoded by individual neurons. This novel phenomenon, which we refer to as the presaccadic encoding expansion, was not present in the single-cell responses and relied on high-dimensional neural representations. This expansion was observed at the level of whole population of simultaneously recorded neurons (using SVM) or even when we examined the interaction between only two neurons (using joint discriminability).

Our results dovetail with recent findings showing the importance of high-dimensional neural representations for different cognitive functions and demonstrate that neural populations can encode information not evident in the firing rate of individual neurons [3,11–13]. Interestingly, we found increased dimensionality for encoding the nonpreferred than the preferred target locations, corresponding with the larger population-level encoding benefit for nonpreferred locations. This boost indicates that a high-dimensional representation is particularly useful for improving population encoding when signal improvement cannot occur through single-cell mechanisms (e.g. gain modulation), which is compatible with a recent finding that reduced dimensionality accompanied poorer performance [3]. In addition, we found that encoding expansion was accompanied by reduction in noise correlation for nonpreferred locations only, indicating the importance of inter-neuronal interactions between neurons selective to different locations. Altogether, our findings suggest that cognitive states modify the information content of prefrontal ensemble activity more easily than that of single-cell activity because of many components that contribute to the population code. For example, the complex manifold of population neural activity could greatly change if the activity of a few neurons is slightly altered by cognitive states.

Here we used two very different measures (decoding accuracy based on the SVM and joint discriminability based on the LDA) to assess the information content of the population response. Importantly, both measures captured changes in the representation of targets between stimulus encoding and saccade preparation. However, only joint discriminability allowed measurement of the information content of pairs of neurons for which signal and noise correlations could be computed as well. Numerous studies have linked reductions in noise correlation to enhanced cognitive functions such as attention [25,26,36]. Here, we used the LDA to isolate the effect of changes in inter-neuronal correlations and found that a presaccadic reduction in noise correlation enhances joint discriminability. Results based on both the SVM and LDA reveal a presaccadic expansion of the population code, with population-level benefits at the nonpreferred locations attributable both to the dimensionality of the population representation and reductions in noise correlation between neurons. Accordingly, we were able to quantify the contributions of both high-dimensional neural representation and reduction in noise correlations within the same study.

Interestingly, a recent study has suggested that the amount of pair-wise correlations could put an upper bound on the dimensionality of neural ensembles [28]. Another study measured the effect of noise correlation on population encoding in prefrontal cortex and found that removing noise correlation increased decoding accuracy for the cue and attention location but not for the saccade [36]. Together, these results suggest that high-dimensional representation and reduction in noise correlation are not necessarily correlated and instead, their relationship depends on cognitive states. Here, we found that both a decrease in NC and an increase in dimensionality contribute to enhanced encoding indicating that the observed presaccadic expansion in the population of FEF neurons occurs due to modulation by an external source (e.g. attentional modulation) as well as changes in interactions between neurons in the same population.

We find that during saccade preparation, the FEF's representation of space undergoes expansion at the level of the population code, showing another example of how a population of neuron can be more informative than individual neurons. A recent study has shown that the FEF neurons are able to encode targets of eye movements far from their classic receptive field [37]. Our results illustrate how population-level changes can further expand the neural code beyond what individual neurons can achieve. This encoding expansion relies on both high-dimensional neural representations and reductions in noise correlation between neurons. Overall, our results demonstrate the ability of prefrontal neural ensembles to recruit non-informative individual neurons to encode a larger part of space, and to exploit high-dimensional neural representations and changes in inter-neuronal relationships for improving the encoding capacity of the whole population according to cognitive states.

## Material and methods

### Experimental paradigm and recording

Two monkeys (macaca mulatta) were trained to perform an oculomotor delayed response (ODR) task. Each trial started with the monkey fixating on the fixation point, followed by the presentation of a visual target (black circle) in one of 16 locations defined with two possible eccentricities (7 and 14 visual degrees) and eight different angles (0, 45, 90, 135, 180, 225, 270, and 315 degrees). The target remained on screen for one second, followed by 1 second of delay after which the monkey could saccade to the remembered location. The monkey was rewarded with a drop of juice if the saccade fell within 5 degrees of visual angle from the target location. The neural recording was done using a V-probe linear array electrode, allowing us to simultaneously record from 16 channels. Using a strict criterion for spike sorting, we identified 188 single-unit (1165 pairs of neurons) and 320 clusters of multi-unit activity (total 508) during the 20 recording sessions (with an average of 784 trials per session). All experimental procedures were in accordance with the National Institutes of Health Guide for the Care and Use of Laboratory Animals, the Society for Neuroscience Guidelines and Policies, and the Montana State Animal Care and Use Committee.

### Data analysis

The baseline mutual information (MI) was computed using neural firing rates during non-overlapping 150 msec windows during the fixation interval for each pair of locations.

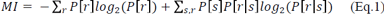

where *s* is the set of stimuli, *r* is the set of neural response (mean firing rate), *P*[*r|s*] is the conditional probability of neural response *r* given stimulus *s*, and *p*[*s*] and *p*[*r*] are the prior probability of stimulus *s* and response *r*, respectively. We refer to this mutual information as 'spatial discriminability' because it quantifies the ability of single neuron to discriminate between various target locations.

Neural selectivity across different epochs was then quantified by comparing the MI for each pair of locations during sensory encoding (50-200 msec time window aligned to the target onset), memory (1500-1650 msec time window aligned to the target onset), and saccade preparation (-120-30 msec time window aligned to the saccade onset), to the baseline MI (using two-sided Wilcoxon ranksum and *p* < .05 as the significant value).

To extract decoding information, we trained a SVM classifier with a linear kernel using the population and single neural responses to 70% of the stimuli. Decoding accuracy was computed by the performance of the classifier on the remaining 30% of the stimuli. We repeated the calculations 500 times and in each repetition, the stimuli were randomly partitioned into training and test sets. In order to compute the dimensionality of neural representation, we first sorted the simultaneously recorded units in each session based on their mutual information (spatial discriminability) and subsequently constructed a set of neural ensembles by subsequently adding new units (with decreasing mutual information). We next computed the overall decoding accuracy as a function of the number of units (NU) in different epochs of the experiment. The dimensionality was defined as the number of units needed to exceed 75% of maximum decoding accuracy. To measure the standard error of the dimensionality, we used a bootstrapping method that sampled the session's ensembles with replacement (1000 times). More details are provided in the Supplementary methods (S1 Text).

The preferred location is defined based on the maximum of the average response (neural tuning curve) or the average decoding accuracy (decoding tuning curves) of locations in visual encoding, memory, and saccade preparation intervals for individual units and the ensemble. For the dimensionality analysis we used an average of the three intervals across all dimensions. We computed the tuning dispersion of spatial tuning by fitting a Gaussian probability density function and using the standard deviation of the fitted function.

Moreover, in order to estimate the information content of a pair of neurons, we applied Fisher linear discriminant analysis (LDA) on each pair of neurons (2-dimensional neural space) and extracted the optimal weight vector *w_opt_* as:

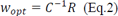

where *R* is the difference between the mean vectors of a given two conditions, and *C*^−1^ is the inverse of the summation of the covariance matrixes of two conditions in the 2-dimensional neural space constructed by a pair of neurons. The joint discriminability was computed by a projection of the 2 locations represented in the 2-dimensional neural space on to the weight vector. The joint discriminability is the mutual information between the projections of the neural response and target location. The geometric mean of MI values in single cells of a given pair was considered as the single-cell version of the joint discriminability, which we refer to as 'isolated' discriminability and was used as a control for measuring encoding expansion (see below).

Different encoding expansion indices were computed using the performance based on the SVM or discriminability as follows:

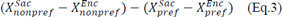

where *X* can be SVM_individual_, SVM_ensemble_, discriminability_isolated,_ or discriminability_joint_. To compute the final measure the encoding expansion index using the SVM analysis or the LDA, we subtracted single-unit expansion index (*X* = SVM_individual_ or discriminability_isolated_) from the corresponding ensemble expansion index (*X* = SVM_ensemble_ or discriminability_joint_).

Statistical analysis were performed using the Wilcoxon's signed-rank test (for paired comparisons) or ranksum (for unpaired comparisons), unless otherwise specified. All *p* values are reported up to three digits and values below 0.001 are all reported as *p* < *10^−3^*.

More details are provided in the Supplementary materials and methods (S1 Text).

## Acknowledgments

We thank Bard Duchaine and Matt van der Meer for helpful comments and Kelsey Clark for helpful comments and edits on the manuscript. The work was supported by MSU startup fund, Whitehall 2014-5-18, NSF BCS143221, and NEI R01EY026924 to BN, Dartmouth startup fund to AS, and by EPSCoR (RII Track-2 FEC) to BN and AS.

## Supplementary figures and figure legends

**Supplementary Figure 1.**
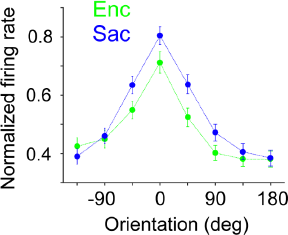
Normalized response of FEF neurons with mixed selectivity to targets presented at different angles relative to their preferred location (0 degree) during visual encoding (Enc, green) and saccade preparation (Sac, blue). The error bars represent s.e.m. Overall, the response profile of neurons with mixed selectivity was very similar to the average of all recorded neurons.

**Supplementary Figure 2.**
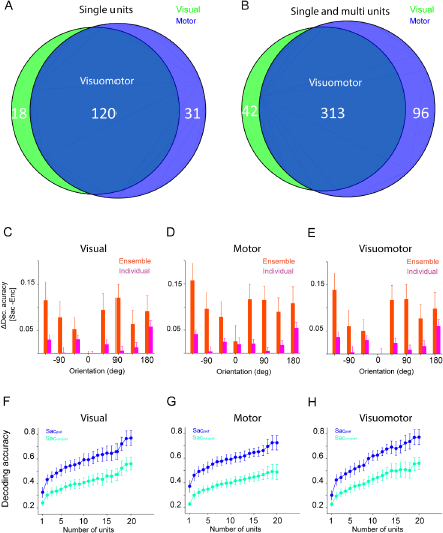
Effect of cell types on expansion and dimensionality of neural representation. Conventions are the same as in Figure 2.

**Supplementary Figure 3.**
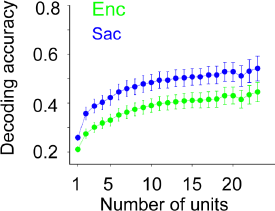
Dimensionality analysis based on the decoding accuracy of the SVM classifier. Plotted is the decoding accuracy of the ensemble of neurons as a function of the number of neurons used for the SVM classifier, separately for visual encoding (Enc, green) and saccade preparation (Sac, blue) epochs. The error bars represent s.e.m.

**Supplementary Figure 4.**
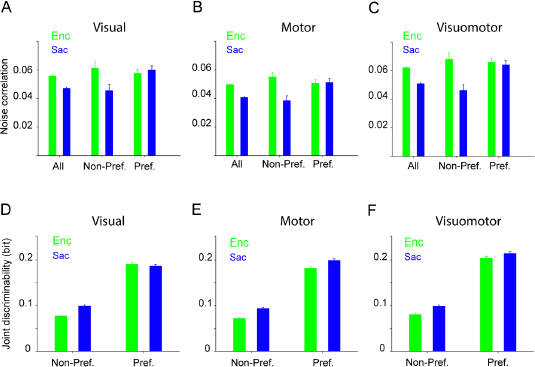
Effects of cell type on noise correlation and joint discriminability. Conventions are the same as in Figure 2.

**Supplementary Figure 5.**
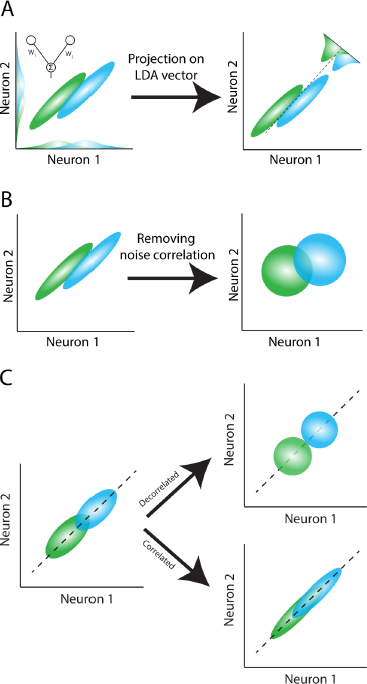
(**A**) Schematic of the linear discriminant analysis (LDA) used for computing joint discriminability. Ellipses represent the variability of neural population response in two conditions (green and blue). The LDA weight vector for the pair of neurons (solid line in the left panel) is the optimum weight vector for population decoding, and is equal to a linear combination of the two neurons that maximally separates neural responses in the two conditions. The dashed line corresponds to the best criterion to discriminate responses to the two conditions (i.e. the LDA line). (**B**) Schematic of the computations of signal and noise correlations. The signal correlation was computed after shuffling the order of repeated trials within each condition (right panel). The noise correlation was obtained by subtracting the signal correlation from the total correlation (i.e. correlation coefficient without shuffling). (**C**) Effects of changes in inter-neuronal correlations on the discriminability of paired neural responses. Plotted is an idealized response of two neurons with positive signal and noise correlations to the target at two different locations (green and blue), and ellipses represent the variability of the neural response. A decrease in the noise correlation between the two neurons (decorrelated case) produces a reduction in the overlap of the distributions of the neural responses to the two stimuli, corresponding to improved discrimination between two locations. In contrast, an increase in correlation between the two neurons (correlated case) creates more overlap between the neural responses to the two stimuli, indicating a drop in discriminability.

**Supplementary Figure 6.**
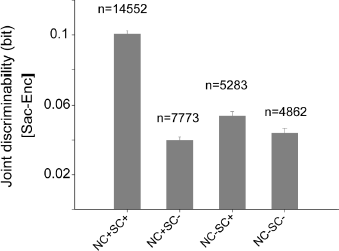
Neuron pairs with positive signal and noise correlations comprised about half of the population, and exhibited a change in joint discriminability twice other pairs of neurons. Plotted is the change in joint discriminability from visual encoding to saccade preparation for various groups of pairs, sorted based on the sign of the signal and noise correlations for the neuron-condition pairs. The number of neuron-condition pairs in each group is shown above each bar.

**Supplementary Figure 7.**
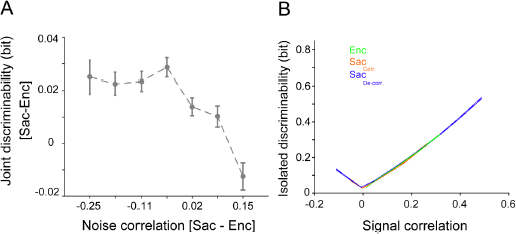
Although the change in the information content of pairs of neurons depends on the change in noise correlation, the information content of individual neurons does not depend on change in noise correlation between visual encoding and saccade preparation. (**A**) Plotted is the change in joint discriminability for pairs of neurons that show a negligible change in SC between visual encoding and saccade preparation (ΔSC < 0.05, *n* = 6419), illustrating the direct contribution of NC changes to joint discriminability. Improved joint discriminability during saccade preparation compared to visual encoding is most prominent when NC is reduced in saccade preparation relative to visual encoding in the same pair. (**B**) Plotted is isolated discriminability as a function of signal correlation computed separately for neuron-location pairs during visual encoding (Enc, green) and during saccade preparation. The latter is plotted for pairs of neurons that showed a large (> 0.1) reduction (blue) or increase (red) in noise correlation from visual encoding to saccade preparation. The shaded area represents s.e.m. This figure provides a control of Figure 4A, and illustrates that the information content of individual neurons in the same pairs of neurons is the same for visual encoding and saccade preparation, and for pairs which become more correlated and decorrelated during saccade preparation.

## Supplementary materials and methods

### General and surgical procedures

two male rhesus monkeys (Maccaca mulatta) were used in this experiment. All experimental procedures were in accordance with the National Institutes of Health Guide for the Care and Use of Laboratory Animals, the Society for Neuroscience Guidelines and Policies, and the Montana State Animal Care and Use Committee. Each animal was surgically implanted with a head post and recording chamber. Surgery was conducted using aseptic techniques under general anesthesia (isoflurane), and analgesics were provided during postsurgical recovery. Eye position monitoring was performed via optical tracking with a high frequency camera (the eyeLink system; 2000 Hz). Eye monitoring, stimulus presentation, data acquisition, and behavioral monitoring were controlled by the Monkey Logic system [1]. Visual targets (black circles with 1 visual angle radius) presented in one of 16 locations defined with 2 possible radii (7 and 14 visual angles) and 8 different angles (0, 45, 90, 135, 180, 225, 270, and 315 degree). Stimuli are presented on an LED-lit monitor (ASUS VG248QE: 24in, resolution 1920x1080, refresh rate 144 Hz) positioned 28.5 cm in front of the animal's eyes.

### Single-neuron recording in FEF

Single-neuron recordings in awake monkeys were made through a surgically implanted cubic titanium chamber (30×30 mm). The sixteen-site, linear V-probe electrode (125 µm distance of sites) was lowered into the cortex using a hydraulic microdrive (Narashige) to record the extracellular activity of single cells. During each experiment, a recording site in the FEF was first localized by the ability to evoke fixed-vector, saccadic eye movements with stimulation at currents of<50µA [2].

### Spike sorting

To isolate single cells' activity, we used multivariate t-distribution decomposition and Expectation-Maximization for clustering [3]. After that, to find the well isolated clusters in spike sorting, we trained a binary classifier for each pair of clusters in one session of recording. For each cluster, we assigned the minimum performance of binary classifiers which separated that cluster from other clusters as an index to measure the isolation power of cluster. For each classifier, the number of samples was equalized and 10-fold cross validation was applied on equalized samples.

### Individual and population neural response

At each time period we quantified the firing rate of individual single or multi-unit clusters. The population activity included all of the simultaneously recorded units in a single recording session. So the population representation of location used for the SVM analysis is a point in R^N^ space, where N is the number of simultaneously recorded units (single and multi) in one session of recording. For neuron-condition pairs, 2 locations were represented in R^2^ neural space.

### Neural response tuning curves

In order to compute the neural response relative to each neuron's preferred location, for each neuron the 8 stimulus locations were circularly shifted such that the zero-degree location indicated the maximum response, computed by averaging the neural response across the three task periods (visual, memory, and saccade).

### Mutual information (MI)

We used mutual information as a measure to quantify the ability of neural responses to represent the target location [4]. Mutual information quantifies how well an ideal observer can discriminate between target locations based on the neural response. We computed mutual information using the neural firing rate during non-overlapping 150 msec windows within different intervals:

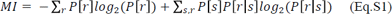

where *s* is the set of stimuli, *r* is the set of neural response (mean firing rate), *P*[*r|s*] is the conditional probability of neural response *r* given stimulus *s*, and *p*[*s*] and *p*[*r*] are the prior probability of stimulus *s* and response *r*, respectively.

### The support vector machine (SVM) classifier

We trained an SVM classifier with a linear kernel (Cortes and Vapnik 1995) using the population and individual neurons responses on a randomly selected 70% of trials as a training set. Then we measured the classification accuracy of the trained classifier on the remaining 30% of trials. Training was done using a least square method for finding the separating hyper-plane [6]. To calculate the standard error of classification accuracy in one session, we repeated the calculations 500 times with different sets of training and test trials. In each repetition, the data were randomly partitioned into training and test sets. We trained and tested the SVM with stimuli represented either in 1 dimension (for 'individual' decoding accuracy), or n dimensions, where n is the number of units in a given recording session (for 'ensemble' decoding accuracy).

### Decoding tuning curves

To compute decoding tuning curves (Fig. 1E) we extracted the classification accuracy of condition C (C is one angle) by using the classifier confusion matrix. The eight possible target locations were circularly shifted such that zero location indicted the location with maximum performance, computed by averaging classification performance over the three task epochs (visual encoding, working memory, and saccade preparation).

### Signal and noise correlation

The signal and noise correlation between each pair of simultaneously recorded units was computed for each pair of target locations (a neuron-condition pair). The correlation coefficient across trials captured the total correlation for each neuron-condition pair. The signal correlation was the correlation coefficient computed after shuffling the order of repeated trials for each target location (Fig. S3B). We repeated the shuffling and calculation of the correlation coefficient 500 times, and used the mean of these *r*-values as the measure of the signal correlation for that neuron-condition pair. The noise correlation was defined by subtracting the signal correlation from the total correlation [7,8].

In Figure 3B, we computed the signal and noise correlations for pairs of conditions for preferred and nonpreferred target locations based on the SVM classifier tuning curves. First we calculated the SVM performance and decoding tuning curve for each session; the target location with maximal decoding accuracy was considered the 'preferred' location, and the location with the lowest decoding accuracy was considered the 'nonpreferred' location. Using these locations, we computed the noise and signal correlations for all pairs of neurons recorded during that session.

### Fisher linear discriminant analysis (LDA) and joint discriminability

In each neuron-condition pair, 2 locations were represented in a 2 dimensional neural space, based on the simultaneously recorded activity of two neurons. We used Fisher linear discriminant analysis (LDA) to find the optimum vector for the linear discrimination between the two responses [9]. For a population of 2 neurons, let *R_s_* (s = 1, 2) as a 2×N matrix (N is number of trials) and 2×2 matrix C_s_ denote the neuronal response covariance matrix for stimulus *s*. Let the 2×1 matrix 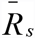 be the average population response to stimulus s. We calculated the optimal weight vector w_opt_ that yields maximum discrimination between two target locations as

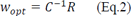

Where *R* is the difference of mean vectors of two conditions 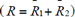 and *C^−1^* is the inverse of summation of covariance matrixes of the two conditions (*C* = *C*_1_ + *C*_2_). After finding *w_opt_*, we projected the population response of each trial onto the optimal weight vector *w_opt_*. This projection is a mapping from two dimensions to scalar. So we could estimate the distribution of projected neural response in one dimension and compute the joint discriminability as the mutual information between the projections of the neural response the target location.

### Dimensionality analysis

We sorted the simultaneously recorded units in each session based on their MI, and made a set of neural ensembles by subsequently adding new units (with decreasing MI). Using this approach, we computed the decoding accuracy as a function of the number of units (NU). Then we fitted 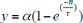 function, where *y* is SVM performance and *n* is number of units, using least-squares and computed dimension as a number of units needed to exceed 75% of maximum decoding accuracy. To measure the standard error of the dimensionality, we used a bootstrapping method that sampled the session's ensembles with replacement (1000 times).

### The tuning curves for noise correlation

To obtain a tuning curve for noise correlation (Fig.3C) we extracted the order of eight possible locations in a manner similar to decoding noise correlation and computed the noise correlation in a given location *C*, by averaging the noise correlation in neuron-location pairs of one session in which pairs were made by locations *C* and its nearby locations (*C*±45 deg).

